# Estimating *de novo* mutation rates using parent-offspring pairs

**DOI:** 10.64898/2026.09.11.751022

**Authors:** Thuy-Trang Nguyen, Matthew W. Hahn

## Abstract

Existing pedigree approaches to identifying *de novo* mutations (DNMs) require at least two parents and a single offspring, limiting applicability. Here, we introduce OOPS (Only One Parent Sequencing), a framework for detecting DNMs using only a single parent-offspring pair. OOPS uses short-read data from the parent and both short and long-read data from the offspring to reconstruct haplotypes in the child, one of which can then be assigned to the sequenced parent. We show that candidate *de novo* mutations from the assigned haplotype can be identified, allowing for estimation of the mutation rate. To demonstrate the accuracy of OOPS, we apply it to a human pedigree in which mutations have also been identified using standard trio-based approaches. OOPS achieves comparable accuracy to trio-based pipelines and recovers consistent mutation rate estimates. By removing the requirement for complete trio sequencing, OOPS expands mutation rate estimation to a wider range of settings.

## Introduction

*De novo* mutations (DNMs) are the source of genetic variation in both evolution and disease (Veltman and Brunner 2012). The per-generation mutation rate is a fundamental parameter in molecular evolution, shaping estimates of divergence times, effective population sizes, and the genetic load carried by populations (Yoder and Tiley 2021). Accurate estimation of the mutation rate is therefore critical across disciplines.

The standard approach to DNM detection in longer-lived organisms requires sequencing a complete trio — both parents and a child — so that variants present in the child but absent from the parents can be identified as *de novo* (Bergeron et al. 2022). Trio-based studies have revealed key features of the mutational process, including a strong paternal bias (Kong et al. 2012; Jonsson et al. 2017; Wang et al. 2020; Wu et al. 2020; de Manuel et al. 2022; Wang et al. 2022a, 2022b; Bergeron et al. 2023; Peña-Garcia et al. 2025; Wooldridge et al. 2025), more minor maternal age-effects (Goldmann et al. 2016; Jonsson et al. 2017; Wang et al. 2025), and inter-individual variation in mutation rates (Sasani et al. 2019). Multi-generation pedigrees have further refined rate estimates, particularly since they can be used to verify Mendelian inheritance of DNMs across generations (Jonsson et al. 2017; Thomas et al. 2018; Porubsky et al. 2025).

While trio sequencing is accurate and can detect mutations transmitted by both parents, it necessarily excludes many species and study systems where two parents cannot be sampled or identified. For instance, in animals with uniparental care, one parent is typically unavailable or difficult to identify. In plants—especially wind-pollinated plants—pollen donors may be harder to identify, even when maternal and offspring tissue can be readily collected together (e.g. acorns attached to oak trees). Even in human clinical studies, one parent is sometimes unavailable or deceased. These constraints have left germline mutation rates unmeasured in many systems.

Recent advances in sequencing technology produce much longer reads, enabling haplotype-resolved genome sequencing of single individuals (Mahmoud et al. 2025). These advances allow for new approaches in mutational studies, overcoming the requirement that complete trios be sequenced. Here, we introduce OOPS (Only One Parent Sequencing), a computational method that estimates the DNM rate from a single parent–child pair. OOPS exploits the ability of long reads to phase the child’s diploid genome into two haplotypes, comparing each haplotype against the single sequenced parent to identify DNM candidates. We validate the approach on a well-characterized human pedigree with high-quality DNM calls (Porubsky et al. 2025).

### Novel Approaches

#### Mutation identification from parent-offspring pairs

OOPS requires both short-read and long-read data from the child, but only short-read data from the single parent (Fig. 1A). As direct input to the program, OOPS requires genotypes from the child and parent (as VCF files), and mapped short and long reads from the appropriate individuals (as BAM files). The pipeline proceeds in three stages (Supplementary Fig. S1).

**Figure 1.**
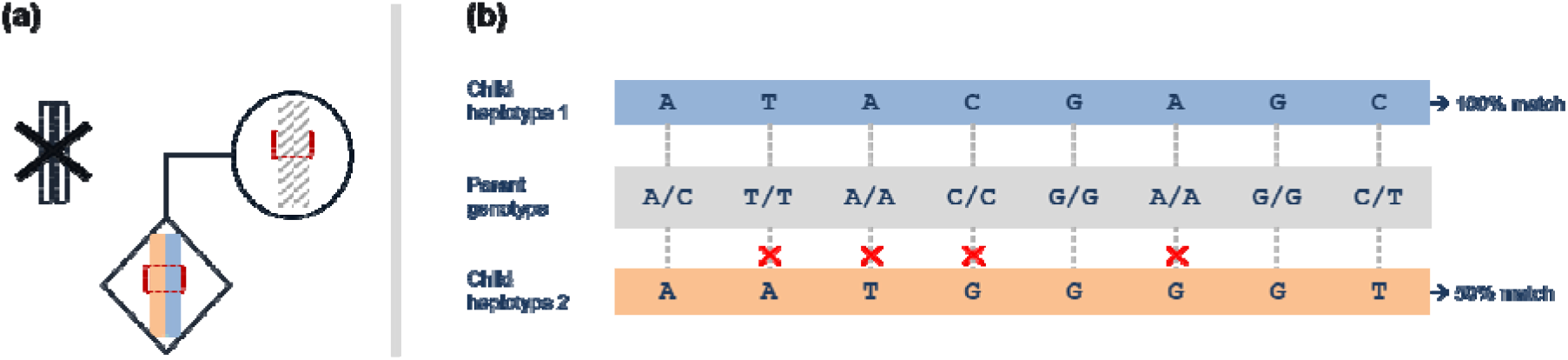
Overview of haplotype assignment and *de novo* mutation detection in the OOPS framework. **(a)** In a pedigree, only one parent and a child are sequenced. Long-read sequencing of the child enables phasing, resolving the diploid genome into two distinct haplotypes (colored blocks). The available parent is sequenced with short reads, yielding unphased genotypes (hatched pattern). **(b)** Within a haplotype phase set (outlined by a red square in panel **a**), the child’s two haplotypes are compared against the parent’s genotype. Here, haplotype 1 (blue) matches the parent at all 8 sites (100%), whereas haplotype 2 (orange) matches at only 4 of 8 sites (50%; mismatches marked with red crosses). This asymmetry implies that haplotype 1 is inherited from the sequenced parent, and the complementary haplotype is inherited from the absent parent, enabling parent-of-origin assignment of haplotypes and candidate *de novo* mutations.

In the first stage, the child’s genotype is phased into haplotype blocks using long-read sequences. The OOPS package includes WhatsHap (Martin et al. 2016) for phasing, but, if preferred, users also have the option of inputting a phased VCF for the child using other software. Most importantly, the child’s diploid genome is now partitioned into phase sets — individual blocks of paired haplotypes (e.g. Fig. 1).

In the second stage, within each phase set, OOPS attempts to assign one of the two child haplotypes to the sequenced parent. As each child haplotype only has one allele, but the parental diploid genotype has two, OOPS scores as a “match” any position in the haplotype that could have come from the parent, and as a “mismatch” any position that could not have come from the parent (Fig. 1B). The expectation is that one child haplotype will match the sequenced parent at nearly every site because it is inherited from that parent; the other haplotype will show many mismatches, though the exact number and proportion depend on many factors. The OOPS default settings for numbers of matches and mismatches needed to assign haplotypes confidently are described in the Supplementary Materials.

OOPS identifies blocks showing the expected asymmetric mismatch pattern: many mismatches on one haplotype and 0 or 1 mismatches on the other. A single mismatch on the low-mismatch haplotype — a site where the child carries an allele absent from the sequenced parent, on the haplotype otherwise concordant with that parent — is flagged as a DNM candidate.

In the third stage, OOPS takes candidate DNMs through three validation steps. First, standard short-read filters are applied (Bergeron et al. 2022), including genotype quality and read-depth in both individuals, homozygosity in the parent, and allelic balance in the child (Supplementary Methods). Second, DNMs must have long-read support: the alternate (mutant) allele must be present on exactly one haplotype in the child, with a minimum number of supporting reads, and a minimum total depth of long-reads. We have found that this step eliminates many post-zygotic mutations often identified by trio-based approaches (e.g. Porubsky et al. 2025). Third, OOPS locally rephases the candidate DNM. WhatsHap is re-run in a 40 kilobase window (adjustable) around each candidate DNM to produce a new phase set, removing false positives caused by haplotype switch errors that accumulate over longer genomic distances. Candidates that pass all three filters are promoted to the final DNM call set.

#### Estimation of the mutation rate

The calculation of a mutation rate requires not just accurate DNM calls, but also the number of sites examined at which a mutation could have been identified—the “callable genome size”— and the false negative rate (FNR), the rate at which true mutations are missed (Bergeron et al. 2022). Using these numbers, the mutation rate within OOPS is calculated as:

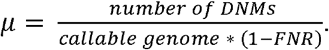

We define the callable genome in OOPS as the total number of bases within phased sets where one child’s haplotype can be unambiguously assigned to the sequenced parent, i.e. phase sets showing the expected asymmetric mismatch pattern (with either 0 or 1 mismatch on the assigned haplotype). To estimate the false negative rate, OOPS samples one high-quality heterozygous SNPs from each assigned phase set when an eligible site is available (Supplementary Materials). The alternate allele at each sampled site is treated as a positive control mutant allele, and is subjected to the same short, long-read validation filters as the DNM candidates, except for the requirement that the sequenced parent be homozygous reference and contain no read support for the alternative allele. The FNR is the fraction of sampled control sites that failed these validation filters.

## Results

### OOPS accurately estimates the mutation rate

We evaluated OOPS using parent-child pairs from the human pedigree sequenced by Porubsky et al. (2025) using multiple sequencing technologies. We focus on a single child (individual NA12879) and analyzed either her mother (NA12878) or father (NA12877) as the single sequenced parent. For this child, Porubsky et al. (2025) reported 6 maternal and 37 paternal autosomal germline single-nucleotide DNMs using a trio-based method.

At 10X PacBio HiFi long-read coverage (down-sampled from the full 37X coverage dataset), WhatsHap produced 17,191 phase blocks with a median length of 42.7 kb, phasing 98.9% of heterozygous sites (Supplementary Table 1). Using the full Illumina short-read dataset for the child and mother (both at 31X coverage), OOPS recovered 2 of the 6 germline DNMs (33%), with 0 false positives (Table 1). In the child-father pair, OOPS made 14 calls: 12 matched the 37 paternal germline DNMs reported by Porubsky et al. (2025), one matched a DNM called in their dataset, but that was unassigned, and one was not in their dataset (i.e. a false positive; Table 1). The majority of the DNMs from the original study missed by OOPS occurred on phase sets for which both haplotypes had many mismatches to the sequenced parent, which appears mainly due to phase-switch errors in haplotyping (see below).

**Table 1.** OOPS performance at 10X long-read coverage on mutations in offspring NA12879. Trio DNMs and rates come from Porubsky et al. (2025). FP = false positives. Mutation rates are expressed per bp per generation.

| Platform | Parent | Trio DNMs | OOPS DNMs | FP | Trio rate | OOPS rate |
| --- | --- | --- | --- | --- | --- | --- |
| PacBio HiFi | Maternal | 6 | 2 | 0 | $0.23 \times 10^{-8}$ | $0.22 \times 10^{-8}$ |
| PacBio HiFi | Paternal | 37 | 14 | 1 | $1.39 \times 10^{-8}$ | $1.53 \times 10^{-8}$ |
| ONT | Maternal | 6 | 1 | 0 | $0.23 \times 10^{-8}$ | $0.25 \times 10^{-8}$ |
| ONT | Paternal | 37 | 4 | 0 | $1.39 \times 10^{-8}$ | $1.02 \times 10^{-8}$ |

Despite recovering only a subset of DNMs in each parent-offspring pair, the mutation rate estimated by OOPS closely matched the trio-based estimates (Table 1). The maternal rate estimated was 0.22 × 10^−8^ per bp per generation (trio-based: 0.23 × 10^−8^) and the paternal rate was 1.53 × 10^−8^ (trio-based: 1.39 × 10^−8^), reproducing the male-biased mutation observed in humans (Kong et al. 2012).

### Effect of long-read technology and read-depth

To determine the effects of long-read technology, we repeated the OOPS analyses using Oxford Nanopore (ONT) reads down-sampled to 10X coverage in the child. Again, 98.9% of the heterozygous sites were phased, but the phase-set structure is different from that obtained with PacBio: WhatsHap produced 440 phase sets with a median length of 690 kb (Supplementary Table 1). In this dataset, OOPS recovered fewer DNMs from both pairs (1 maternal, 4 paternal) and produced a slightly underestimated paternal mutation rate (Table 1). The smaller number of DNMs detected using the ONT data seems to be due to frequent phase-switch errors, with many long haplotypes showing regions with low numbers of mismatches and regions of high mismatches (Supplementary Figure S2). In these cases, OOPS cannot confidently assign haplotypes.

To determine the effects of long-read depth in the child, we analyzed PacBio HiFi datasets with 5X, 7X, 10X (the analysis above), 15X, and the full 37X coverage. Paternal DNM calls increased steadily with depth, while maternal calls plateaued (Fig. 2a; Supplementary Table 2). False positives were rare at every depth tested: one was called at 5X in the child-mother pair and one at 10X in the child-father pair, and none at any other depth. The estimated mutation rate remained stable across read-depths (Fig. 2b; Supplementary Table 2), as both the callable genome and FNR were also recalculated at each read-depth.

**Fig 2.**
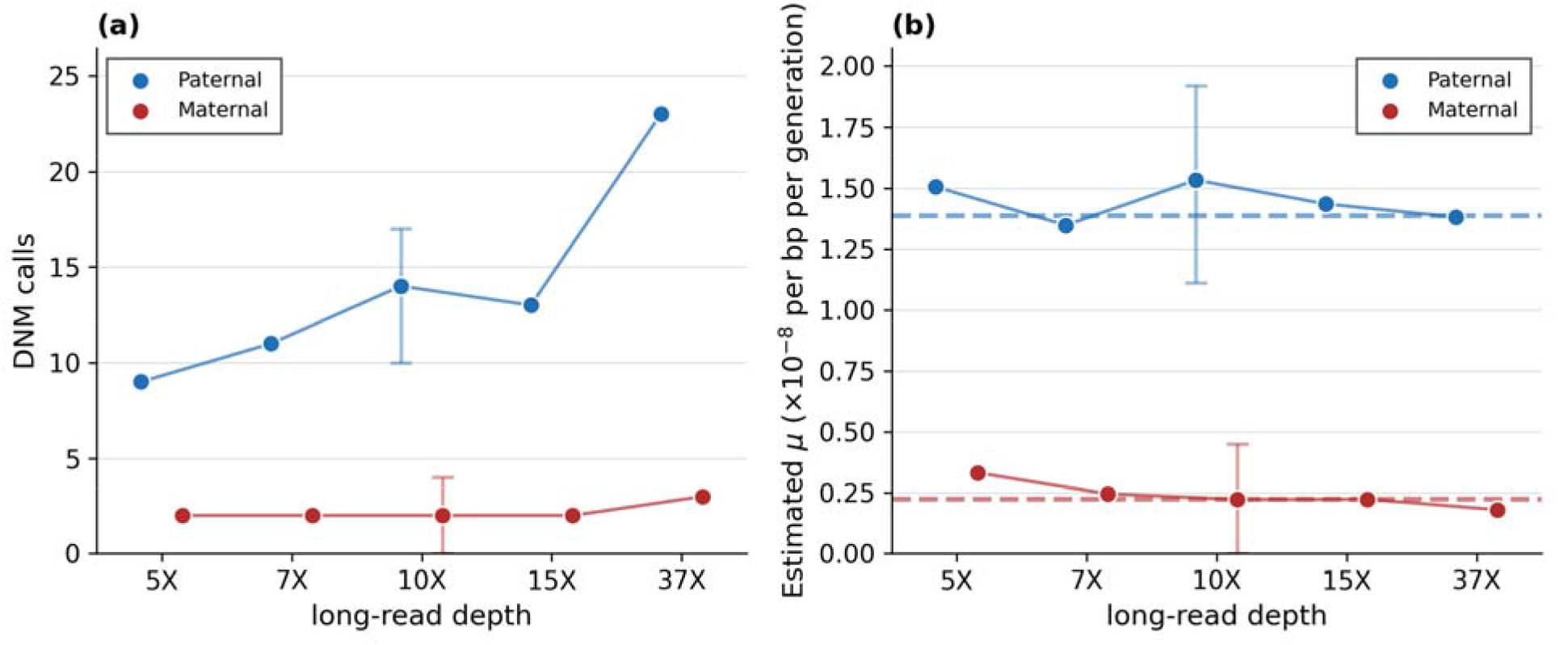
*De novo* mutation detection as a function of PacBio HiFi read depth. (**a**) Number of DNM calls of paternal (blue) and maternal (red) origin recovered by OOPS at coverage levels from 5X to 37X. (**b**) Estimated per-generation mutation rate at each depth, with the trio-based rates of Porubsky et al. (2025) shown as dashed lines. Bars at 10X give the 95% confidence interval across 100 independent samplings of the same data — of the number of calls in (**a**) and of the estimated rate in (**b**) (see also Supplementary Table 3).

Because there is stochasticity associated with read-sampling at every depth, some variation in coverage-specific results could arise from read sampling itself. We quantified this variability by repeating the 10X HiFi analysis 100 times, each time drawing a new set of reads and rerunning phasing and all OOPS steps. Across replicates, the mean number of calls was 13.8 for the paternal pair (range 9–19) and 2.2 for the maternal pair (range 0–4) (Fig. 2a; Supplementary Table 3). The mean mutation-rate estimates were 1.54 × 10^−^□ for the paternal pair and 0.25 × 10^−^□for the maternal pair; the corresponding 95% confidence intervals are shown in Figure 2b (see also Supplementary Table 3). These replicates capture variability arising from read-sampling, phasing, and candidate filtering at 10X coverage, encompassing the range of mutation rate results captured across different read-depths (Fig. 2b).

## Discussion

We demonstrate here that accurate estimation of the per-generation mutation rate is possible with sequencing data from only a parent-child pair. The key insight is that long-reads can phase a child’s genome into haplotypes, one of which can often be accurately assigned to the sequenced parent. Although OOPS recovers only a minority of true DNMs — because only sites within well-phased blocks with sufficient read support are assessable — the mutation rate estimate remains accurate because the callable-genome calculation corrects for the reduced available genome (see also Figure S5 in Wang et al. 2020).

The method has several limitations. First, accuracy depends critically on phasing quality: haplotype switch errors can cause false positives by placing a true inherited variant on the wrong haplotype and can cause false negatives by making it difficult to assign haplotypes to the sequenced parent. We have found that the local rephasing step partially mitigates this issue, at least by eliminating many false positives. The same general issue is also why we have chosen not to use phased data from the sequenced parent: in addition to false positives caused by allelic gene conversion (cf. Narasimhan et al. 2017), switch errors or true recombination events in the parent will lead to more noise but not necessarily more detectable mutations. See Boukas et al. (2026) for a similar method that does use a phased parent, and Young et al. (2024) for a method using a pair of phased siblings.

As a second limitation, OOPS is currently restricted to single nucleotide variants on autosomes: indels, structural variants, multinucleotide mutations, and mutations on sex chromosomes are not detected. However, all of these restrictions can be relaxed in the future. Any type of mutation can be included, including structural variants identified using the longreads themselves. With slight modifications, OOPS can be run on sex chromosomes, a natural application for chromosomes that are often hemizygous and that can therefore be phased even without long-reads.

Despite its limitations, OOPS fills an important gap in evolutionary genomics. Germline mutation rates have been measured using trio sequencing in a limited number of species, predominantly those that can be raised in the lab or a zoo (Bergeron et al. 2023). OOPS enables DNM-rate estimation in many non-ideal circumstances, with relatively low requirements for read-depth. As long-read sequencing costs continue to decline, this approach could dramatically expand the taxonomic breadth of mutation-rate estimates.

## Acknowledgements

We thank Richard Wang, Yadira Peña-García and Jeff Rogers for valuable feedback. This work was supported by NIH grant R01-HD107120.

## Data Availability

The pipeline is available at https://github.com/TrangNg-Th/OOPS-Only-One-Parent-Sequencing and is distributed as a Conda package (oops-dnm) for straightforward installation.

