## Supplementary tables and figures for "Estimating *de novo* mutation rates using parent-offspring pairs"

### Materials

**Data.** We used publicly available sequencing data from Porubsky et al. 2025 (CEPH 1463). The focal pairs included: NA12879 (child), NA12878 (mother), and NA12877 (father). PacBio HiFi and ONT long reads for the child, Illumina short reads for the parents, as well as the Illumina short-read genotypes for the child and parents were obtained via the AWS Open Data program (s3://platinum-pedigree-data/data/) as well as the European Nucleotide Archive (BioProject: [PRJEB86317](https://www.ebi.ac.uk/ena/data/view/PRJEB86317)). PacBio HiFi reads were down-sampled to 5X, 7X, 10X, and 15X using samtools (Li et al. 2009). All coordinates are from the T2T-CHM13v2.0 reference genome (Nurk et al. 2022), stored at: <https://s3-us-west-2.amazonaws.com/human-pangenomics/T2T/CHM13/assemblies/analysis_set/chm13v2.0_maskedY_rCRS.fa.gz>

**Phasing and haplotype assignment.** Phasing was performed with WhatsHap v1.7 (Martin et al. 2016). The child genotypes from the joint VCF file (i.e. the file including the parental genotypes as well) were phased using the child's long-read BAM, producing phase-set-tagged genotypes. In each phase set, each position present in child and parent was subjected to further filters based on the joint VCF before matches and mismatches were called. These included requiring an Illumina read-depth between 15X and 50X (derived from the average read depth 31X +/- 3$\sigma$ ), a genotype quality of at least 30, and at most two alleles.

After filtering, for each position within each phase set (i.e. haplotype block), OOPS compares the allele on each child haplotype against the parent's genotype. If the child allele is included in one of the two parent alleles, it is considered a match; if not, it is a mismatch. The haplotype–parent mismatch count is then computed as the total number of mismatches in a phase set.

Not every phase set has two haplotypes that can be confidently assigned to a parent of origin. We used three criteria for assigning haplotypes. First, a phase set had to contain at least as many SNPs as the median number of SNPs per phase set in that run. This cutoff is computed per run and therefore differs sharply between platforms: for the PacBio HiFi data at 10X coverage it was 12 SNPs, whereas for the ONT data at 10X coverage it was 3,376 SNPs, because ONT produces far fewer and far longer phase sets (Supplementary Table 1). Second, one haplotype must have either 0 or 1 mismatches—this is inferred to be the transmitted haplotype; a haplotype carrying exactly 1 mismatch is additionally a DNM-candidate block. Third, the two haplotypes must be strongly asymmetric: the ratio of the smaller to the larger mismatch count must be below 0.1. For a haplotype carrying a single mismatch, this means the other haplotype must carry at least 11 mismatches; for a haplotype carrying none, the requirement is satisfied by any non-zero mismatch count on the other haplotype.

**Callable genome estimation.** The callable genome size equals the sum of lengths of all phase sets that pass the above filters, including assigned haplotypes with both 0 and 1 mismatch.

**DNM candidate identification.** Blocks in which one haplotype carried exactly 1 mismatch, and the other showed at least 11 mismatches, were retained as DNM-candidate blocks. Candidate DNMs were further filtered as follows. Using the short reads, the sequenced parent must be homozygous REF at the candidate position, with no reads supporting the mutant (ALT) allele. Candidate sites were further filtered by read-depth (15–50X in both child and parent), genotype quality (≥30) in both child and parent, only two alleles, and >5 reads for both REF and ALT in the child. Candidates falling within homopolymer runs ≥8 bp were discarded. Because per-haplotype mismatch counts are calculated after applying the read-depth and genotype-quality filters, a haplotype recorded as carrying a single mismatch could have contained additional mismatches that were filtered out. To exclude possible mismatch-rich regions of low quality, we therefore required that no additional mismatch on the candidate-bearing haplotype, counted before these filters were applied, occur within 2 kb of the candidate site.

Using long reads, the two alleles in the child were required to be supported by reads assigned to opposite haplotypes: reads supporting one allele had to be assigned exclusively to one haplotype, and reads supporting the other allele exclusively to the other haplotype. Only reads with both base quality ≥20 and mapping quality ≥20 were counted. For the results reported in the main text, we additionally required at least one long read supporting the candidate ALT allele and at least nine long reads covering the site in total.

For candidate DNMs that pass the above filters, we performed local rephasing to improve the accuracy of the parental assignments. For each candidate, we extracted a ±20 kb window around each candidate from the joint VCF (i.e. 40 kb total) and re-ran WhatsHap using long reads. Because these local windows are small and contain fewer variants, we relaxed the two thresholds applied to whole-genome phase sets. First, the variant-count cutoff is set to the 10^th^ percentile rather than the median (for PacBio HiFi data at 10x coverage, this corresponds to requiring only one genotyped site in the newly inferred phase set). Second, the asymmetry threshold was relaxed from 0.1 to 0.5, allowing a haplotype carrying the single candidate mismatch to be retained provided that the other haplotype carries at least 3 mismatches. Candidates whose newly inferred phase sets do not satisfy these relaxed criteria are discarded.

**False negative rate calculation.** To quantify the number of DNMs that could have been missed by our pipeline, we constructed positive-control sites from haplotype blocks confidently assigned to the sequenced parent. Within each assigned block, we identified heterozygous sites in the child for which the ALT allele lay on the haplotype assigned to the sequenced parent. To ensure that these sites represented true variants transmitted from that parent, we further required the sequenced parent to be heterozygous for the same ALT allele. We randomly sampled one such site per assigned block to avoid over-weighting larger blocks. Consequently, the maximum number of sampled sites was determined by the number of assigned blocks available in each run.

Each sampled site was then subjected to the same short-read and long-read validation filters applied to candidate DNMs, except for the requirement that the sequenced parent be homozygous REF and lack support for the candidate ALT allele, because the sampled control sites were intentionally selected to be heterozygous in the parent. The false-negative rate (FNR) was calculated as the fraction of sampled control sites that failed these validation filters.

**Trio-based mutation rate calculation.** The trio-based mutation rate from Porubsky et al. (2025) was calculated as the number of germline DNMs attributed to each parent, divided by the average callable genome size across samples in their study (see Supplementary Table 10 in Porubsky et al. 2025; this number is 2,666,892,426 callable sites). This calculation yields a maternal mutation rate of 0.225 x 10^-8^ per site per generation (6 / 2,666,892,426) and a paternal mutation rate of 1.387 x 10^-8^ per site per generation (37 / 2,666,892,426).

**Software availability.** OOPS is implemented as a Bash/Python pipeline with Slurm job manager and is available under the MIT license at https://github.com/TrangNg-Th/OOPS-Only-One-Parent-Sequencing. A Conda package (recipe/) is provided for dependency management. The pipeline requires bcftools, samtools, tabix, WhatsHap, pysam, pandas, numpy, scipy, and matplotlib.

**Supplementary Figure 1. Detailed pipeline of OOPS.** OOPS uses short- and long-read data from the child and short-read data from one parent, provided as VCF and BAM files. First, the child’s heterozygous variants are phased with WhatsHap, or a user-supplied phased VCF, to partition the genome into paired haplotype blocks (phase sets). Within each phase set, both child haplotypes are compared position by position with the sequenced parent’s diploid genotype; the haplotype showing near-complete concordance is assigned to that parent, whereas the other is expected to contain multiple mismatches. A unique mismatched allele on the otherwise concordant parental haplotype is identified as a candidate DNM. Candidates are evaluated using short-read quality, depth, parental homozygosity, and child allelic-balance filters; long-read haplotype-specific support; and finally, locally rephased in an adjustable 40-kb window to remove false positives caused by haplotype-switch errors. Candidates passing all validation steps are retained in the final call set.

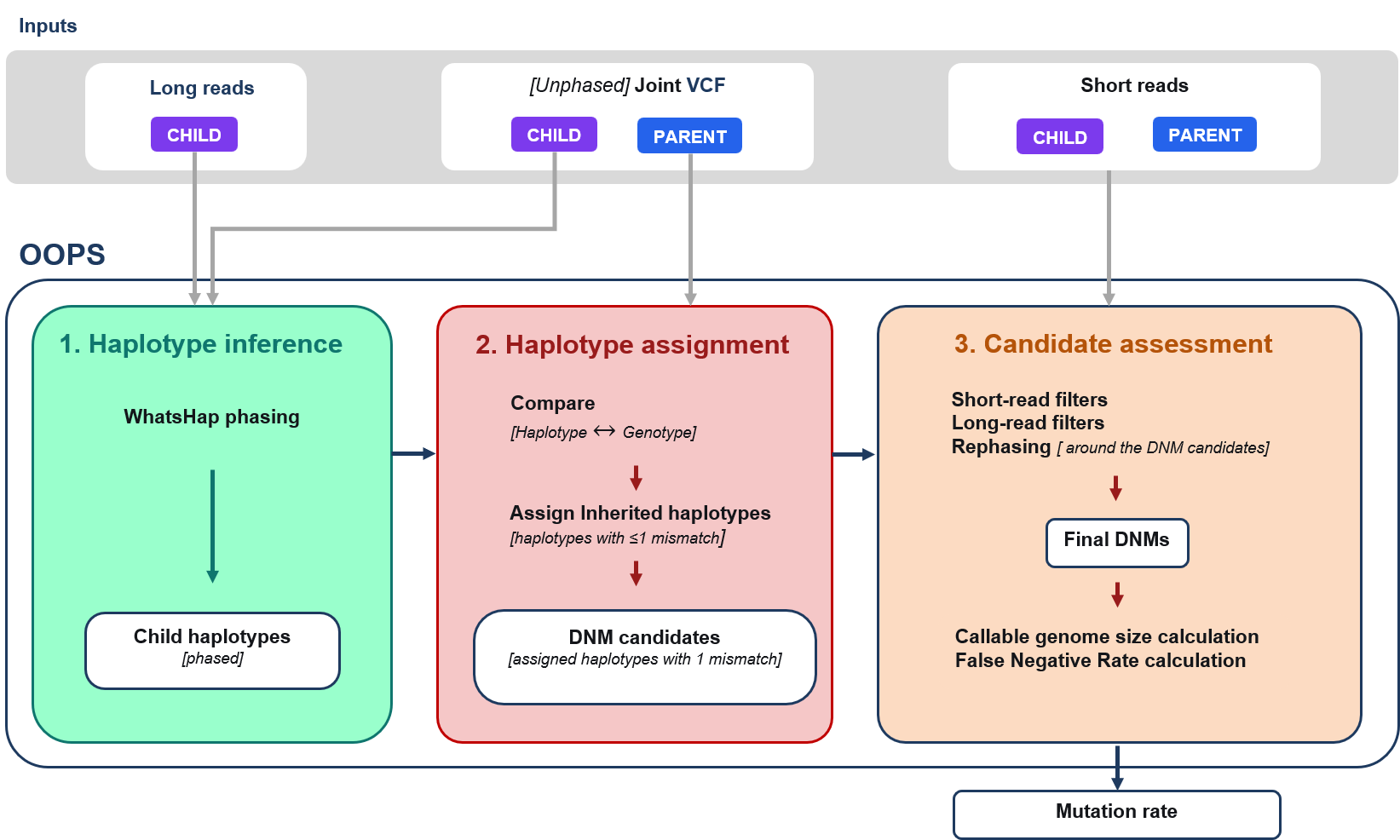

**Supplementary Figure 2. Example of phase-switch errors in an ONT-based phase set.** The child (NA12879) was phased with WhatsHap from ONT reads and compared to the sequenced maternal genotypes (NA12878). Mismatch fractions were calculated in 50-kb sliding windows. For each haplotype, a mismatch fraction of 0 indicates that all variants in the window are consistent with one of the two alleles carried by the parent, whereas a value of 1 indicates that none are consistent. H1 is plotted upward in blue and H2 downward in red on the same 0–1 scale. In the absence of a phase-switch error, the haplotype inherited from the sequenced parent is expected to remain close to zero across the phase set, whereas the other haplotype is expected to have a higher mismatch fraction. The orange dashed line marks the inferred phase-switch position at approximately 104.4 Mb. To the left of the switch, H2 is the parent-matching haplotype, whereas to the right, H1 is the parent-matching haplotype. The switch divides the phase set into two long segments, with the phase-set-wide mismatch counts for H1 and H2 that are both high (H1 = 8,209; H2 = 4,981).
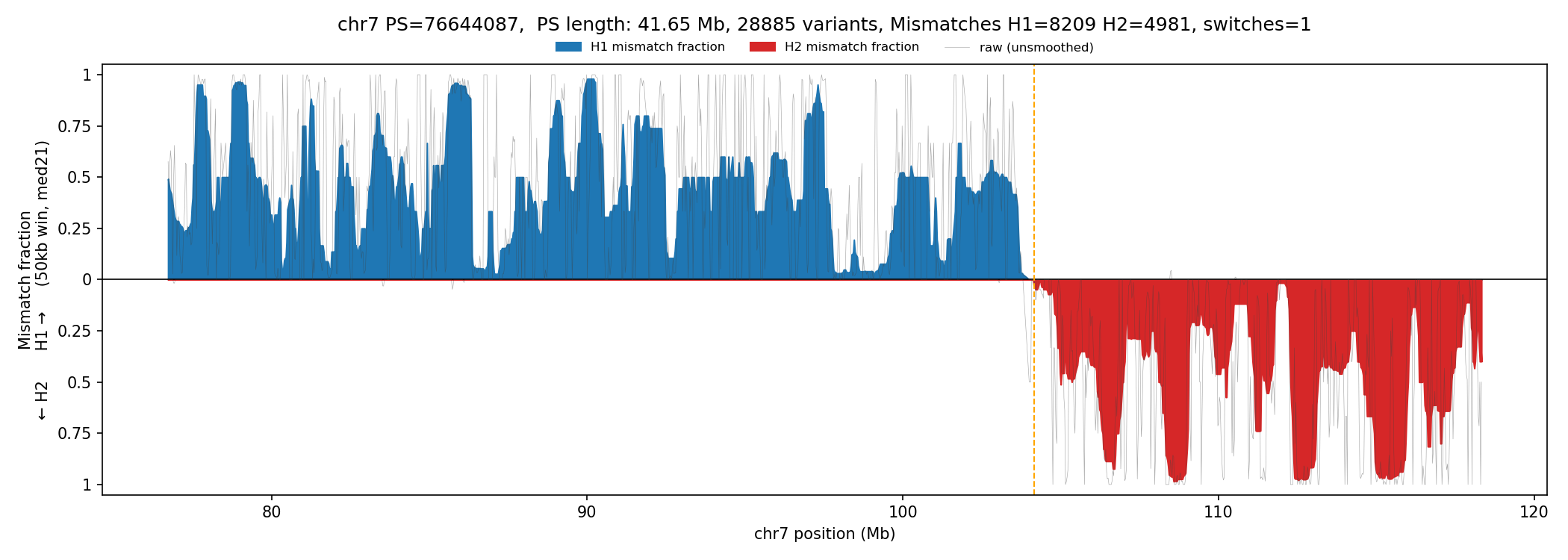

**Supplementary Table 1. Phase-set and SNP counts for offspring NA12879 at 10X PacBio and ONT coverage.** The variant-count cutoff is the median number of variants per phase set and is calculated within each run.

|  | **PacBio HiFi 10X** | **ONT 10X** |
| --- | --- | --- |
| Phase sets | 17,191 | 440 |
| Median phase-set length (kb) | 42.7 | 689.7 |
| Median SNPs per phase set (IQR) | 13 (4-162) | 262 (11–7,439) |
| Phase sets scored by OOPS, after site-level filters | 14,367 | 247 |
| Median SNPs per scored phase set (after filtering) | 12 | 3,376 |

**Supplementary Table 2. OOPS results across PacBio HiFi sequencing depths in offspring NA12879.** A DNM not found in Porubsky et al. (2025) is counted as a false positive (FP).

| **Depth** | **Paternal Calls** | **Maternal calls** | **Paternal FP** | **Maternal FP** | **Paternal rate** | **Maternal rate** |
| --- | --- | --- | --- | --- | --- | --- |
| 5X | 9 | 2 | 0 | 1 | 1.51 x 10^-8^ | 0.33 x 10^-8^ |
| 7X | 11 | 2 | 0 | 0 | 1.35 x 10^-8^ | 0.25 x 10^-8^ |
| 10X | 14 | 2 | 1 | 0 | 1.53 x 10^-8^ | 0.22 x 10^-8^ |
| 15X | 13 | 2 | 0 | 0 | 1.44 x 10^-8^ | 0.22 x 10^-8^ |
| 37X | 23 | 3 | 0 | 0 | 1.38 x 10^-8^ | 0.18 x 10^-8^ |

**Supplementary Table 3. Variability of the OOPS estimate across 100 independent samples of the PacBio HiFi data for offspring NA12879 to 10X coverage.** Replicates were run with different random seeds, with each replicate representing a separate random draw of reads, phased and analyzed from scratch.

|  | **Paternal (NA12877)** | **Maternal (NA12878)** |
| --- | --- | --- |
| Number of replicates | 100 | 100 |
| Average DNM calls (range across replicates) | 13.8 (9–19) | 2.2 (0–4) |
| True positive counts (average) | 13.3 | 1.9 |
| False positive counts (average) | 0.52 | 0.33 |
| Mutation rate, mean [95% confidence interval, × 10⁻⁸] | 1.54 × 10⁻⁸ [1.11, 1.92] | 0.25 × 10⁻⁸ [0.00, 0.45] |
